# A Shared Adenocarcinoma Transcriptomic Program Enables Prediction of Therapeutics Applicable Across Tissues

**DOI:** 10.1101/2025.10.15.682703

**Authors:** Mary Deng, Zhitong Zhou, Christopher E. Lietz

## Abstract

**Background:** Adenocarcinomas are malignancies arising from glandular epithelial cells or secretory tissue, and account for most cancer-related mortalities worldwide, despite advances in treatment. New treatments are often developed in the context of the organ from which the tumor originates. However, there exists shared biology across glandular epithelium at the molecular level regardless of organ system, and there has been a shift towards the molecular classification of tumors. Defining and targeting pan-adenocarcinoma specific molecular features may facilitate the development of treatments applicable across adenocarcinomas, expediting drug discovery efforts and translation to the clinical setting.

**Methods:** We performed an integrated transcriptomics analysis of adenocarcinomas originating from different organ systems. RNA sequencing expression profiles from lung adenocarcinoma (LUAD), stomach adenocarcinoma (STAD), and colorectal adenocarcinoma (COAD) with matched normal tissue from The Cancer Genome Atlas (TCGA) was used to discover a pan-adenocarcinoma specific transcriptomic module. A standardized DESeq2 based pipeline was used to test for differentially expressed genes (DEGs) between each adenocarcinoma and its matched normal tissues, followed by cross-cancer comparisons to identify a consensus transcriptional module. Enrichment analysis was performed to reveal biological pathways associated with the consensus module, with a particular focus on those involved in oncogenesis. Prognostic significance of the consensus transcriptional module was evaluated using survival modeling, and the module was tested against in vitro transcriptomic drug perturbation signatures using Connectivity Map (cMAP) analysis to identify candidate drugs targeting adenocarcinomas, regardless of organ system of origin.

**Results:** Despite the diversity of adenocarcinomas tested, there existed a significant and large overlap of the genes dysregulated across tested tumors. A consensus transcriptomic module was defined, and it predicted patient prognosis in each of the three adenocarcinomas. Leveraging the top shared biomarkers through cMAP analysis, we identified 36 FDA-approved drugs that are capable of reversing the shared malignant transcriptional module towards the normal state. Among the FDA-approved drugs were the EGFR and ALK inhibitors gefitinib and crizotinib, both currently used for treating LUAD, providing validation the pipeline could discover efficacious drugs. Taken together, we developed an approach for organ system independent cancer biomarkers and drug discovery, and leveraged it to identify drug candidates for expanded use in adenocarcinomas. The FDA-approved drugs identified through this pipeline serve as candidates to repurpose as pan-adenocarcinoma anti-cancer therapeutics, reducing both time and cost for drug development.

**Conclusion:** Our research provided insights into the common molecular mechanisms across multiple adenocarcinomas and unveiled potential drug candidates for future therapeutic testing.

## Introduction

Adenocarcinomas are the most common histologic subtype of human cancers such as those arising from lung, colon, pancreatic, and gastric (Bray et al. 2018; Siegel et al. 2023; Sung et al. 2021). Although breast and prostate adenocarcinomas are also highly prevalent, they are largely sex specific. While overall cancer incidence has been decreasing, rates continue to be high and colorectal adenocarcinoma incidence has alarmingly been rising in young people (Sinicrope 2022). Collectively, nearly one third of people in the United States will be diagnosed with an invasive form of these adenocarcinomas in their lifetime. Adenocarcinomas continue to drive significant mortality, with mortality rates for colorectal cancer, non-small cell lung cancer, pancreatic cancer, and gastric cancer at 35.9, 35.0, 11.1, 3.5 deaths per hundred thousand US citizens, respectively (Collaborators 2023; Didier et al. 2024; Kratzer et al. 2024).

There have been meaningful advances in the treatment of these tumors, especially with precision oncology approaches. In colorectal adenocarcinoma, the mutational status of *RAS* or *BRAF* guides applications of targeted inhibitors, and mutational burden informs application of immunotherapy (Biller & Schrag 2021). Non-small cell lung cancer treatment is now informed by actionable genomic alterations in *EGFR, ALK, ROS1, KRAS*, and *NRG1* (Jeon et al. 2025). Patients with pancreatic cancer may benefit from targeted agents depending on *BRCA* mutations or genomic alterations of *NTRK* or *KRAS* (Christenson et al. 2020). Gastric adenocarcinoma treatment is increasingly biomarker-driven, based on microsatellite instability, PD-L1, HER2, tumor mutational burden, and Epstein-Barr virus (Joshi & Badgwell 2021).

These developments have largely occurred independently for each cancer type. However, recent studies have demonstrated shared targetable biology across adenocarcinomas from distinct organ systems (Liu et al. 2018). It has further been shown that histology-driven transcriptional programs are different between adenocarcinomas and squamous carcinomas of the same organs (Lin et al. 2017). In light of these findings, and the fact that developing new cancer drugs is a daunting task both costly and susceptible to failure, with a success rate of less than 4% in clinical trials (Aitken 2023; Sun 2022), an approach to develop drugs based on histologic subtype could accelerate the development for common malignancies, such as adenocarcinomas.

To address this, we performed an integrative cross-comparison analysis of lung adenocarcinoma (LUAD), stomach adenocarcinoma (STAD), and colorectal adenocarcinoma (COAD), three of the most clinically significant adenocarcinomas that occur in both sexes. Using RNA-sequencing data from The Cancer Genome Atlas (TCGA), we systematically identified individual cancer-specific and a consensus transcriptional module across these adenocarcinomas. Furthermore, pan-adenocarcinoma mechanisms of oncogenesis were identified through pathway enrichment studies. The pan-adenocarcinoma transcriptional module was found to predict patient prognosis, indicating its potential as prognostic biomarkers for adenocarcinomas, and potential to stratify patients to alternative therapies. To discover therapies that target shared adenocarcinoma biology, we leveraged the Touchstone database and Connectivity Map (cMAP) analysis and identified candidate compounds that reverse the shared malignant phenotype (Lamb et al. 2006; Subramanian et al. 2017). Together, our research provided insights into the common molecular mechanisms across multiple adenocarcinomas and unveiled potential drug candidates for future therapeutic testing.

## Material & Methods

### Data acquisition

Datasets for unisex adenocarcinomas of the lung, stomach, and colon were extracted from The Cancer Genome Atlas (TCGA; https://portal.gdc.cancer.gov/) (accessed July 16, 2025), using the [TCGAbiolinks](https://doi.org/10.1093/nar/gkv1507) R package (v.2.36.0) (Figure 1A). All samples and data were obtained through IRB approved projects at other institutions. For each cancer type, we downloaded both gene expression quantification (raw counts) and corresponding clinical metadata, including demographics, overall survival time, and vital status for all patients. Genomic data included the RNA sequencing gene expression of 60,660 genes. Clinical annotations included sample type (primary tumor or matched normal sample controls), patient age at diagnosis, gender, race, and overall survival time and event.

**Figure 1:**
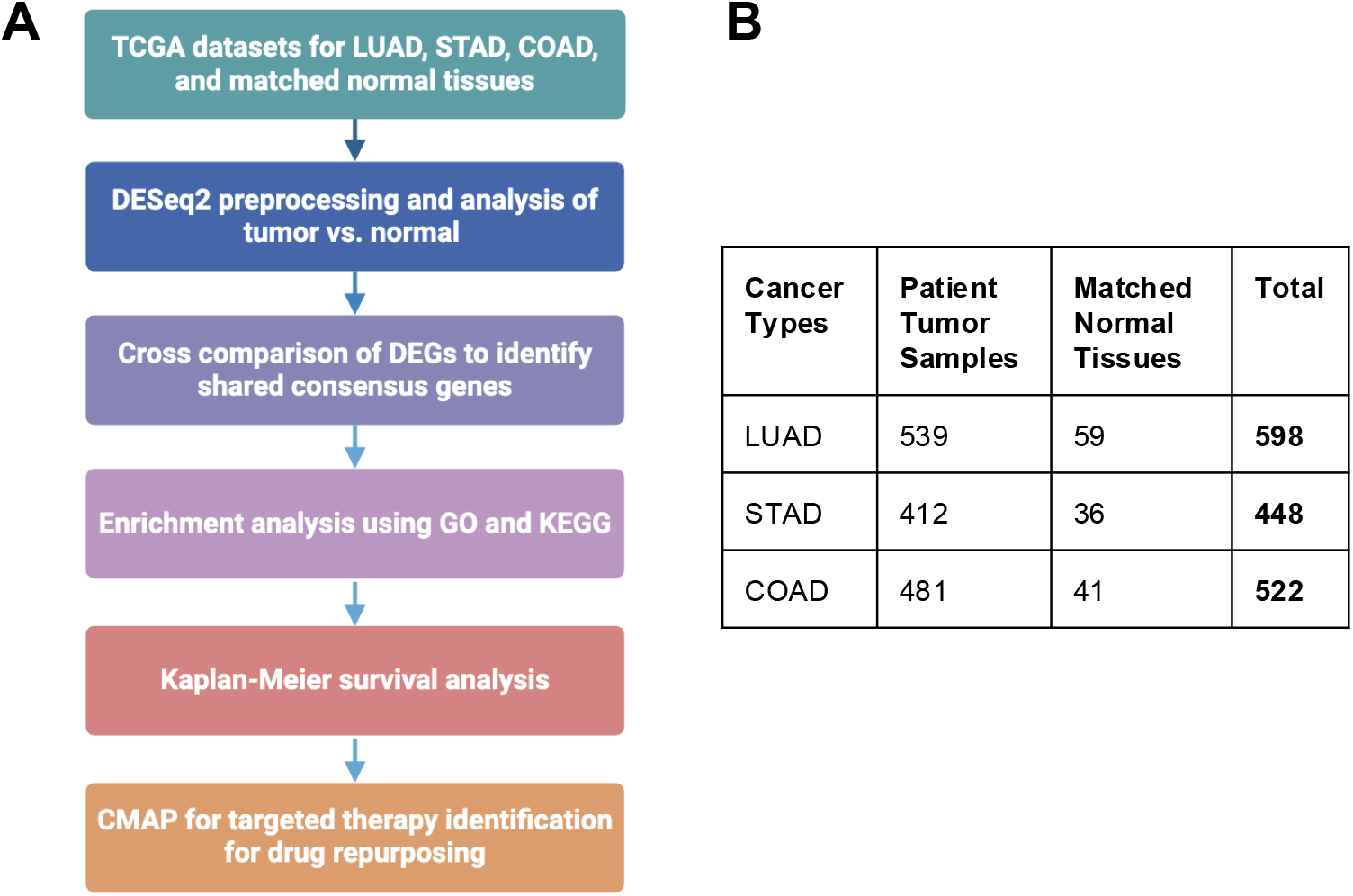
The analysis pipeline and samples acquired. (A) The workflow of the analysis pipeline. (B) Sample number of LUAD, STAD, and COAD, as well as their matched normal tissues acquired from the TCGA.

### Data preprocessing

Samples with clinical annotations (e.g., complete survival data) and RNA sequencing profiles were included in analysis. Samples from metastatic sites were excluded to minimize potential biological variation. The gene expression matrix was filtered to exclude genes with low expression (< 10 counts in two samples or more). In total, the LUAD, STAD, and COAD datasets included 1432 primary tumors and 136 adjacent normal tissue samples (Figure 1B). Data preprocessing was conducted using the R version 4.5.1 and Bioconductor package DESeq2 version 1.48.1 (Love et al. 2014).

### Differential expression analysis

The DESeq2 R package (v1.48.1) was used for differential gene expression analysis after normalizing the raw counts to the size factor of each sample and transforming them into log_2_ space (Love et al. 2014). Differentially expressed genes (DEGs) for each comparison (LUAD *vs*. normal lung tissues, STAD *vs*. normal stomach tissues, and COAD *vs*. normal colon tissues) were identified based on the following criteria: absolute log2 fold change (|log2FC|) ≥ 1 and adjusted p-value (Benjamini-Hochberg method) < 0.05 (Yoav Benjamini 1995).

The DEGs acquired from each cancer *vs*. normal tissue comparison were then subset into upregulated (log2FoldChange > 1) and downregulated (log2FoldChange < -1) genes. To identify the consensus co-expression gene modules shared across LUAD, STAD, and COAD, we intersected significant DEGs obtained from each adenocarcinoma type.

### Principal component analysis (PCA)

PCA was applied for dimensionality reduction on our RNA-seq matrix, which consists of a filtered subset of gene expression features across all samples. Prior to PCA, we first applied a variance-stabilizing transformation (VST) to the raw counts matrix using the DESeq2 package.

### Gene ontology and pathway enrichment analysis

Gene Ontology (GO) enrichment analysis and Kyoto Encyclopedia of Genes and Genomes (KEGG) pathway analyses were conducted on consensus DEGs using clusterProfiler (org.Hs.eg.db; v3.21.0), and clusterProfiler (v4.16.0) was used to convert Ensembl into Entrez IDs (Wu et al. 2021; Yu et al. 2012).

### Unsupervised hierarchical clustering

Unsupervised hierarchical clustering and heatmap generation was performed using the R package pheatmap (v1.0.13) and gplots (v3.2.0). Clustering was performed using Euclidean distance for rows, correlation distance for column, and the average linkage method. Expression values were z-scored across genes during heatmap generation.

### Survival analysis

Gene signature scores were generated for each patient by calculating the mean normalized expression of all consensus genes. Patients were stratified into high- and low-expression groups using the median or optimal cutpoint, which was determined by the maximally selected rank statistics with the survminer surv_cutpoint function (Schumacher 1992). Kaplan-Meier survival analysis was performed separately for LUAD, STAD, and COAD using the survfit function and statistical significance was determined with the log-rank test (p-value < 0.05) using the ggsurvplot function.

### Targeted therapy screening

Connectivity Map ([cMAP](https://www.broadinstitute.org/connectivity-map-cmap)) was employed to identify candidate drugs (Lamb et al. 2006; Subramanian et al. 2017). Queries were implemented on cMAP’s cloud-based computing infrastructure, cMAP and LINCS Unified Environment (CLUE). Query parameters were set as Transcriptomics L1000, Touchstone-L1000, and Recent.

### Visualization

DEGs were visualized with volcano plots using ggplot2 (v3.5.2) for each comparison between adenocarcinoma (LUAD, STAD, or COAD) and its matched normal tissue. Shared DEGs were visualized using VennDiagrams. We also used UpsetR (v1.4.0) to visualize the overlapping genes across different cancer types. Heatmaps were generated using the pheatmap package (v1.0.13) and gplots package (v3.2.0). MA plots were generated using the DESeq2 package (v1.48.1). Venn diagrams were generated using the VennDiagram package (v1.7.3). Statistically significant GO biological functions for both upregulated and downregulated gene lists were visualized in barplots (P ≤ 0.05 and FDR ≤ 0.05) (Yoav Benjamini 1995). Dot plots were utilized to visualize statistically significant KEGG top-ranked metabolic and signal transduction pathways.

### Statistics

Statistical analyses were performed using R software (v4.3.2). For all statistical comparisons, p-values were adjusted for multiple testing using the Benjamini-Hochberg false discovery rate (FDR). A threshold of FDR < 0.05 was considered statistically significant.

## Results

### Identification of cancer-specific and consensus DEGs in LUAD, STAD, and COAD

To discover transcriptional alterations in adenocarcinomas, we conducted differential gene expression analysis on publicly available RNA-seq data of patient samples from TCGA (Figure 1A). We included unisex adenocarcinomas from multiple organ systems to identify shared biological features. Adenocarcinomas are derived from epithelial gland tissues, thus providing a biological justification for the downstream transcriptomic comparison. Patient samples included both primary tumors (n=1432) and matched adjacent normal tissues (n=136) from LUAD, STAD, and COAD patients (Figure 1B) with RNA sequencing profiles from TCGA (Cancer Genome Atlas 2012; Cancer Genome Atlas Research 2014a; Cancer Genome Atlas Research 2014b).

Dimensional reduction of transcriptional data of all filtered genes was performed with principal component analysis (PCA), and PCA plots showed tumor and normal samples clustered separately for LUAD and COAD samples (Supplementary Figure 1A-C), whereas STAD samples were more intermixed than the other two cancer types but still showed a clear separation. Each transcriptomic dataset was normalized separately, and differentially expressed transcripts (|log_2_FC| > 1, FDR < 0.05) between tumor and matched adjacent normal tissue were identified using DESeq2 (Figure 1) (Love et al. 2014). To examine the normalization methods, we also generated MA (mean expression vs. log fold change) plots for each comparison (Supplementary Figure 2A–C). MA plots demonstrated appropriate normalization, with the point cloud centered near M = 0 across the expression range and a reasonable dispersion that decreased with increasing average abundance. Volcano plots illustrate the magnitude and direction of DEGs in each comparison (Figure 2A–C). Among the three adenocarcinomas, LUAD had the largest number of upregulated genes (n=11,682), while STAD had the smallest number (Figure 2D). The number of downregulated genes varied from 2,955 in LUAD to 3,805 in COAD (Figure 2D).

**Figure 2:**
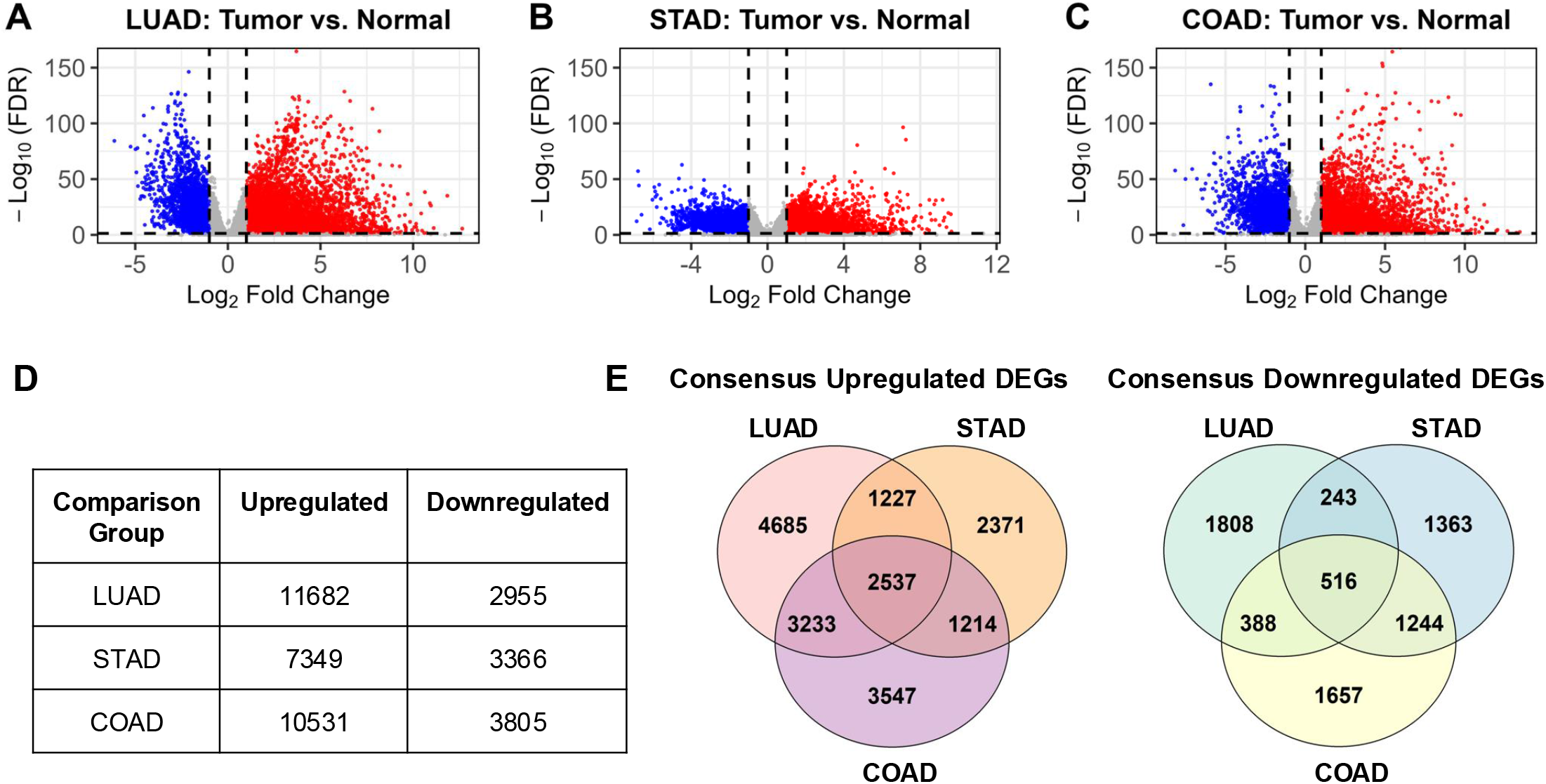
Cancer-specific differential expression genes (DEGs) for each adenocarcinoma vs. its normal tissue comparison and consensus DEGs shared across the three adenocarcinomas. (A-C) Volcano plots of DEGs of LUAD (A), STAD (B), and COAD (C), compared to their respective matched normal tissues. (D) The number of cancer-specific upregulated (red) and downregulated (blue) DEGs in each comparison. (E) Venn diagrams visualizing shared and unique subsets of upregulated (left) and downregulated (right) DEGs.

To test if there exists a conserved transcriptional module across adenocarcinomas, we further analyzed the DEGs in three cancer types through pairwise and triple-wise comparisons (Figure 1A). We found that 2,537 genes were consistently upregulated and 516 genes downregulated across LUAD, STAD, and COAD. The results were displayed using both Venn diagrams and UpSet plots (Figure 2E and Supplementary Figure 3), which provide complementary views of set intersections and shared signatures. The above results show that adenocarcinomas from different tissue of origin share a conserved gene module.

### Oncogenesis and development pathways are enriched in consensus DEGs

To explore if certain biological processes or pathways are enriched in the adenocarcinoma gene module, we conducted Gene Ontology enrichment analysis for consensus gene sets (Gene Ontology Consortium, 2021). Notably, the upregulated genes were significantly enriched in biological processes involving mitosis and proliferation (Figure 3A). “Nuclear division,” “organelle fission,” “nuclear chromosome separation,” and “mitotic cell cycle phase transition” are hallmark processes of proliferation, which are often dysregulated in cancer. Contrarily, consensus-downregulated genes were enriched in tissue-specific and homeostatic functions (Figure 3B). Enriched GO terms included “muscle system process,” “regulation of monoatomic ion transport,” “vascular process in circulatory system,” and “extracellular matrix organization”. Downregulation of these pathways may reflect the loss of tissue identity during tumor progression.

**Figure 3:**
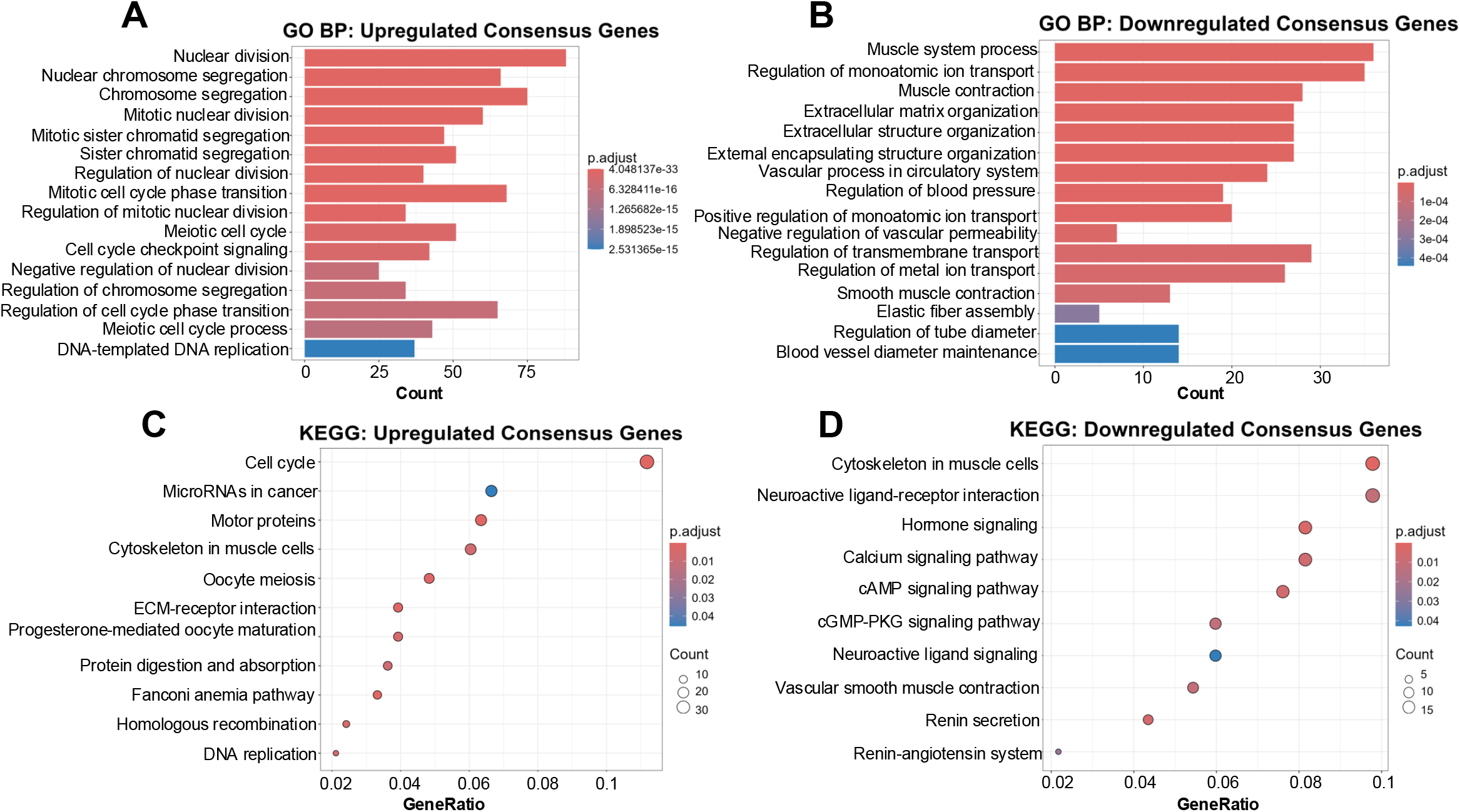
Pathway enrichment analysis reveals the top oncogenic and functional pathways. (A–B) GO enrichment barplots for upregulated and downregulated consensus DEGs shared by all three adenocarcinomas vs. normal tissue comparisons. (C–D) KEGG enrichment dotplots for upregulated and downregulated consensus DEGs by all three adenocarcinomas vs. normal tissue comparisons.

Similarly, KEGG enrichment analysis indicated that consensus-upregulated genes were involved in developmental process and oncogenic signaling pathways, such as “Cell cycle,” “MicroRNAs in cancer,” “Cytoskeleton in muscle cells,” and “Oocyte meiosis” (Figure 3C). Consensus-downregulated genes were enriched in metabolic process and differentiation pathways, including “Neuroactive ligand-receptor interaction,” “Hormone signaling,” “Calcium signaling pathway,” and “cAMP signaling pathway” (Figure 3D). These results were consistent with the transcriptional reprogramming observed in various human cancers.

### Prognostic value of consensus gene modules

To understand if the identified DEGs can serve as biomarkers to predict patient prognosis, we first tested whether the upregulated and downregulated DEGs from all adenocarcinomas are associated with patient overall survival. Kaplan–Meier survival analysis using median expression as the cutoff showed that higher expression of the cancer upregulated DEGs is associated with poor overall survival for each adenocarcinoma, although only significant in LUAD and COAD (Figure 4A-C). Interestingly, the lower expression of the downregulated genes is associated with significantly poor survival only in LUAD patients (Figure 4D), but with better survival in STAD patients (Figure 4E). Similar but more significant results were observed when using the optimal cutoff for patient survival analysis (Supplementary Figure 4).

**Figure 4:**
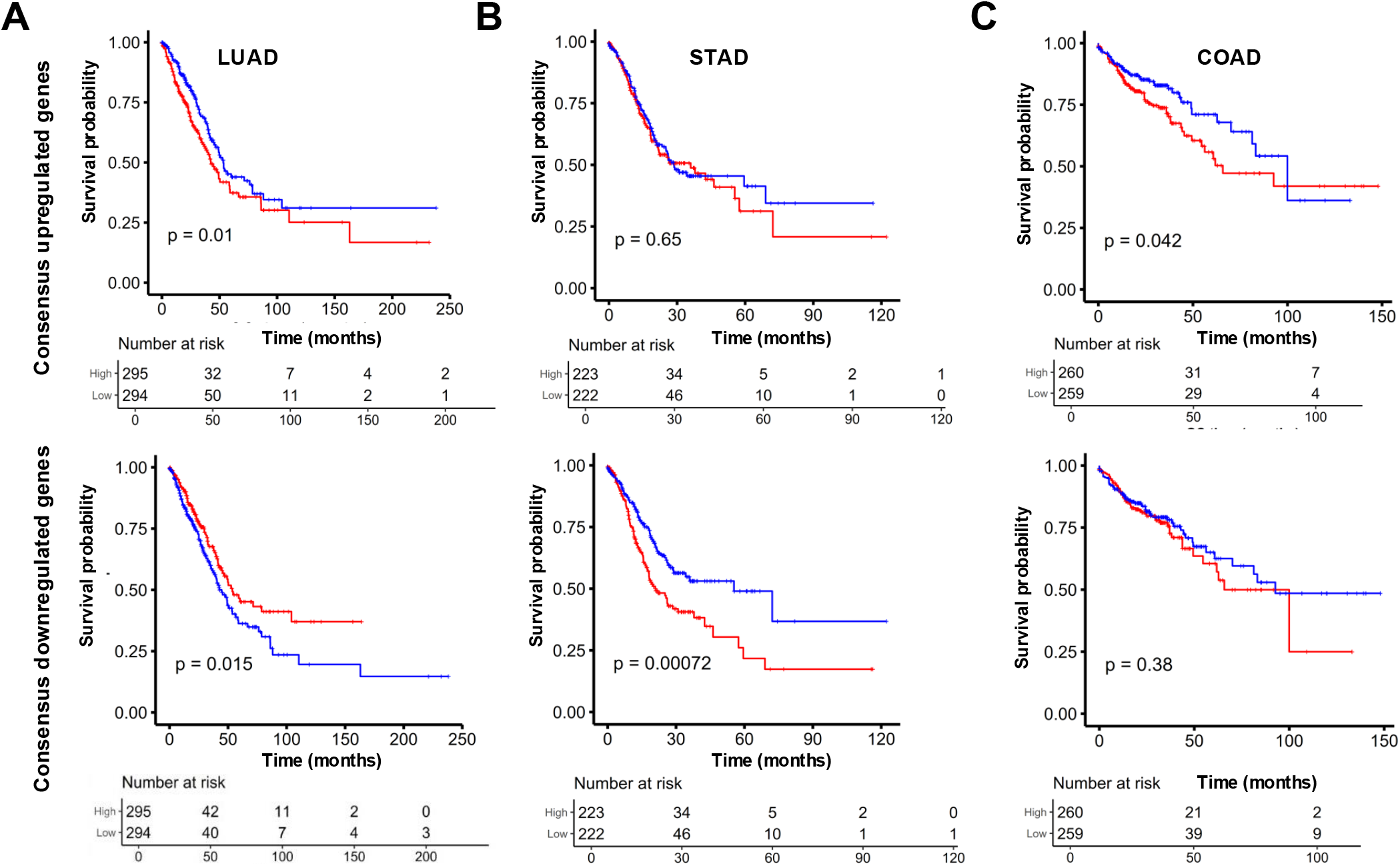
The consensus DEG module can predict the survival of patients with any of the three adenocarcinomas. (A-C) Kaplan–Meier survival analysis of overall survival of patients with LUAD (A), STAD (B), or COAD (C), segregated by the expression of all of the consensus upregulated (top) or downregulated (bottom) DEGs shared by all three adenocarcinomas. Patients were stratified by median cutoff. Statistical analysis was conducted by the log-rank test.

Next, we separated patients into two or three subgroups, based on the consensus gene module through unsupervised hierarchical clustering, and performed Kaplan–Meier survival analysis. Among the three adenocarcinomas, only LUAD patient groups had significantly different survival when both two and three groups were analyzed (Figures 5-6). Despite a trend in survival segregation, STAD and COAD patients from all subgroups had similar survival outcomes (Supplementary Figures 5-8). There were no significant differences in survival for STAD or COAD patients when they were segregated into two and three subgroups (Supplementary Figures 5-8). Taken together, the consensus transcriptional module holds prognostic power for the adenocarcinomas, and, with future refinement, could be used as prognostic biomarkers.

**Figure 5:**
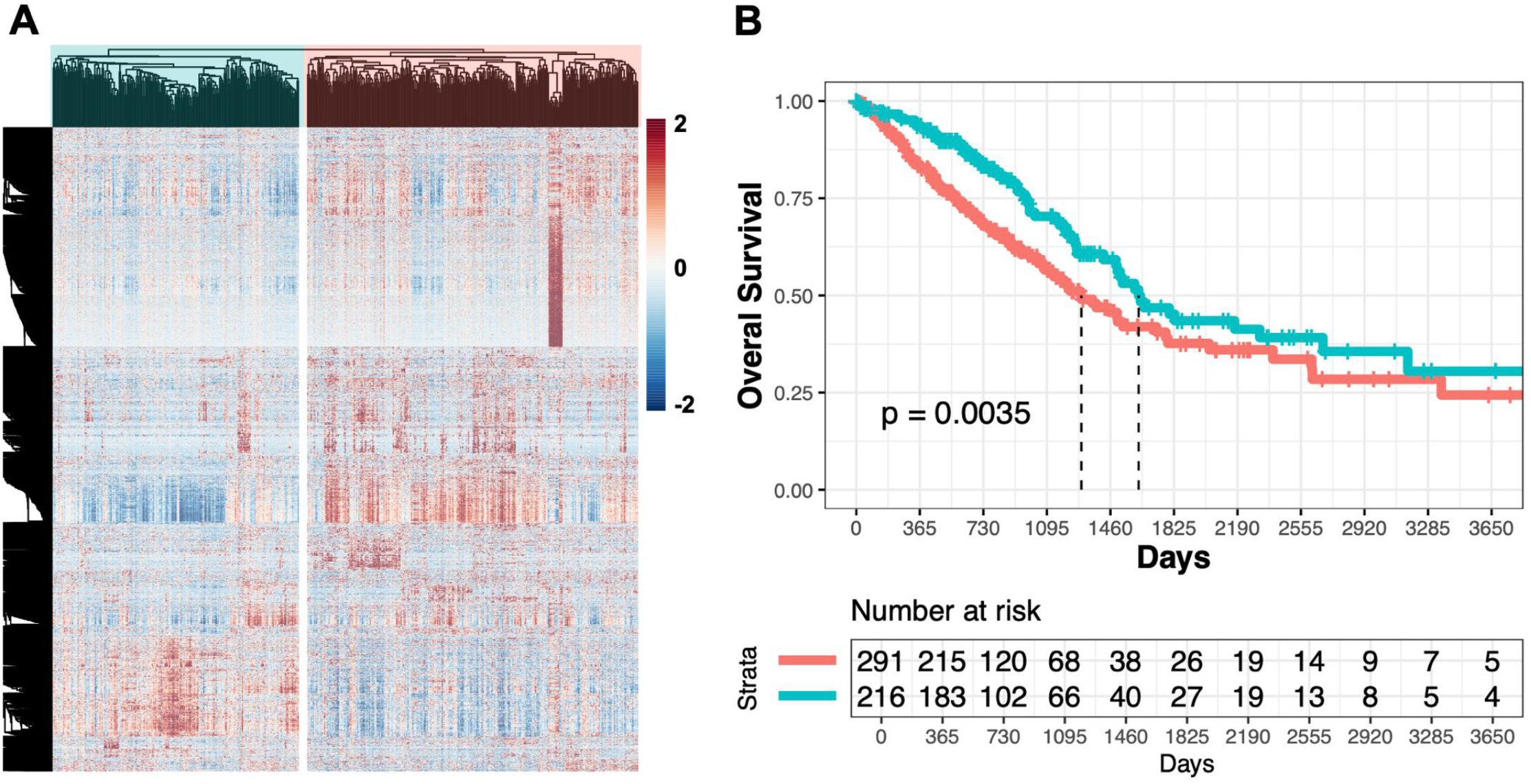
Consensus DEG clusters are associated with LUAD patient survival. (A) Heatmap of unsupervised hierarchical clustering of the 3053 consensus DEGs shared by all three adenocarcinomas categorize LUAD patients into two subgroups. (B) Kaplan-Meier plot of the two primary patient clusters identifies that blue-shaded patient group has better survival than pink-shaded group (p = 0.0035). Statistical analysis was conducted by the log-rank test.

**Figure 6:**
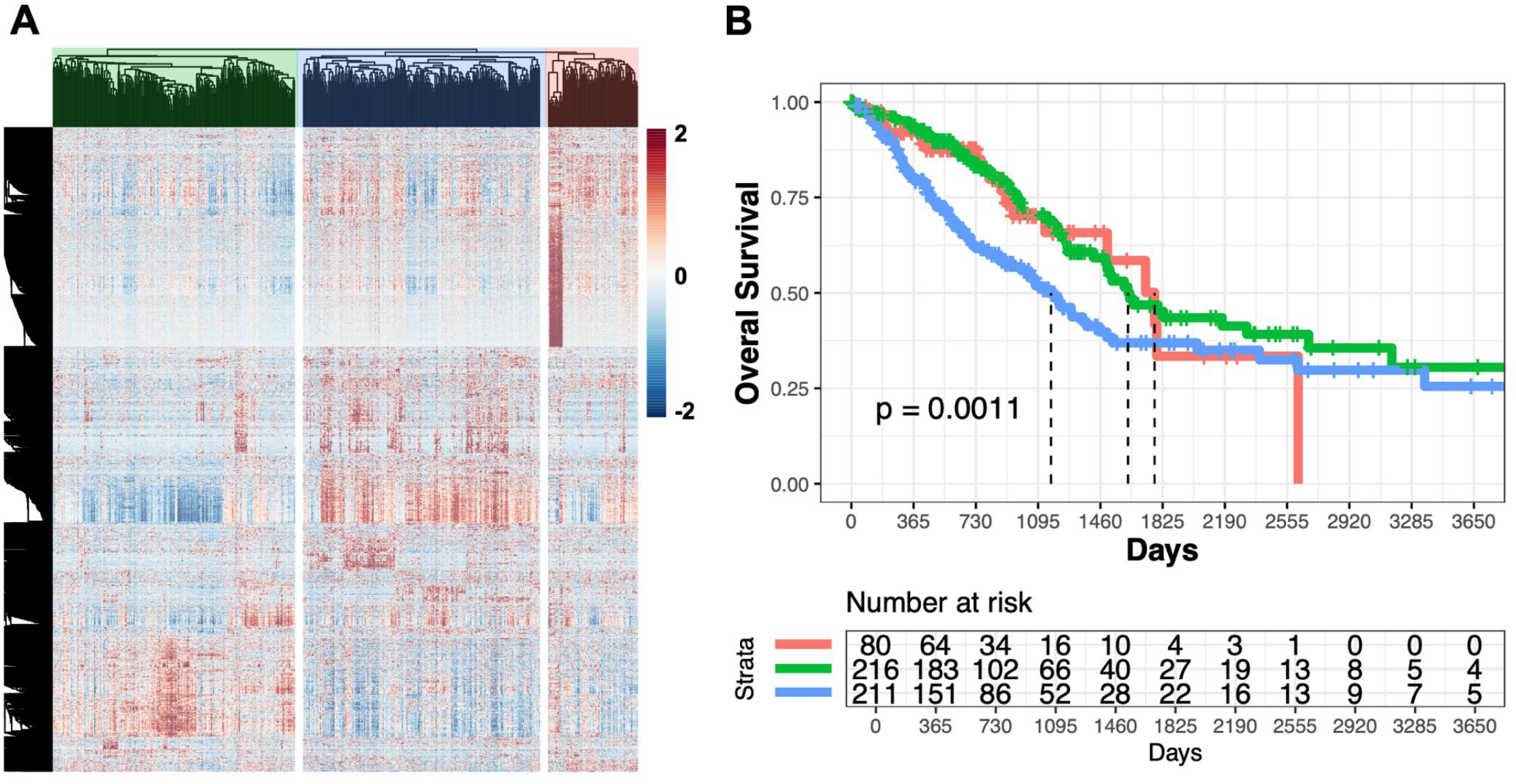
Consensus DEG clusters are associated with LUAD patient survival. (A) Heatmap of unsupervised hierarchical clustering of the 3053 consensus DEGs shared by all three adenocarcinomas categorize LUAD patients into three subgroups. (B) Kaplan-Meier plot of the three primary patient clusters reveals that DEG clusters predict patient survival (p = 0.0011). Statistical analysis was conducted by the log-rank test.

### Utility of identified pan-adenocarcinoma transcriptomic module for novel drug discovery

After identifying the pan-adenocarcinoma transcriptomic module from LUAD, STAD, and COAD, we utilized cMAP analysis to identify drug classes that can revert the expression towards a more normal-like state, reasoning these compounds could be candidates for pan-adenocarcinoma applications. We identified of over 100 classes of small-molecule inhibitors that can reverse the shared malignant transcriptional module towards the normal state (Supplementary Table 1). Among them, there were 36 FDA-approved drugs (Table 1). These small-molecule inhibitors represent drug candidates that can be repurposed to treat multiple adenocarcinomas as well-tolerated, affordable, and viable options. Importantly, among the FDA-approved drugs were the EGFR and ALK inhibitors gefitinib and crizotinib, both currently used for treating LUAD, providing validation the pipeline could discover efficacious drugs. Hence, the pan-adenocarcinoma transcriptomic module enabled us to identify drug candidates that can be potentially repurposed for treating multiple adenocarcinomas.

**Table 1:**
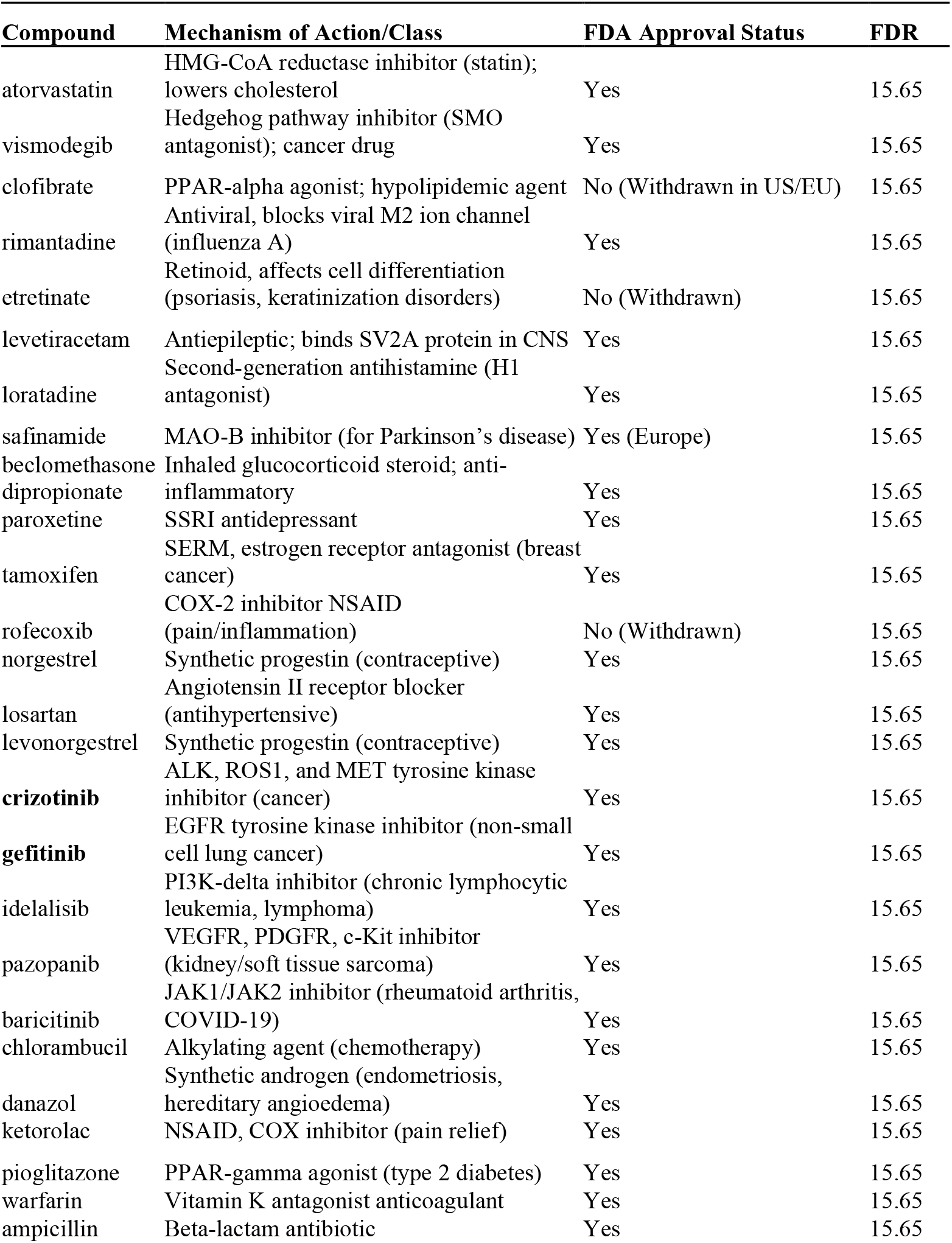
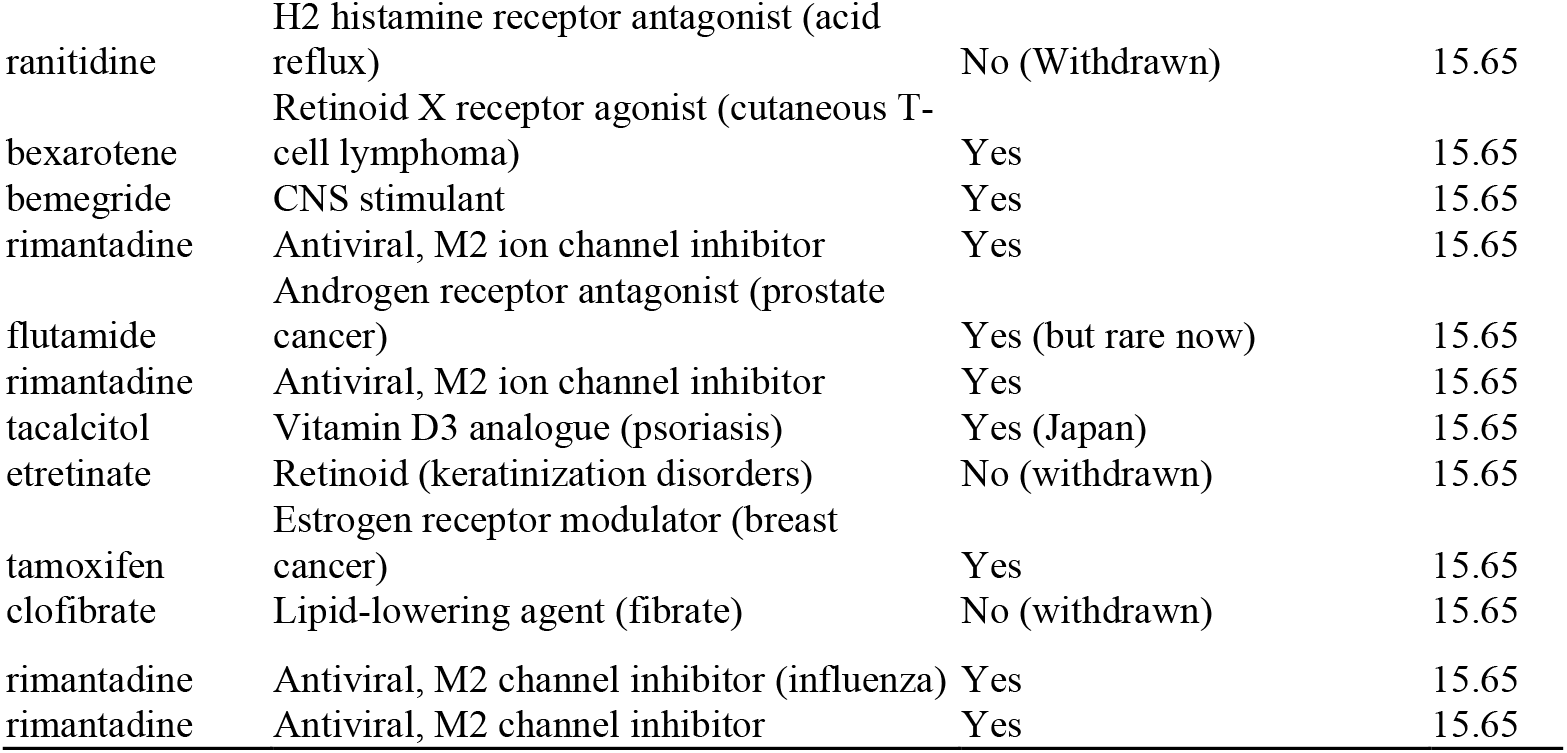
CMAP analysis reveals FDA-approved drugs.

## Discussion

We identified a transcriptomic program consistently dysregulated in lung, gastric, and colorectal adenocarcinomas relative to matched normal tissues. Despite thousands of dysregulated genes in each tumor type, there was a high degree of overlap across adenocarcinomas, supporting the notion they have shared biological features. These transcriptomic features were robust to down-sampling and demonstrated prognostic power across multiple adenocarcinomas. Collectively, this pan-adenocarcinoma transcriptional phenotype highlights molecular commonalities that transcend organ systems and may represent therapeutically actionable biology.

Previous studies have established that tumors from different organs can share molecular features (Hoadley et al. 2018). Lin *et al*. showed that histologic classes such as adenocarcinoma and squamous cell carcinoma explain more transcriptomic variation than tissue of origin (Hoadley et al. 2018). Other analyses have also identified adenocarcinoma subtypes within the gastrointestinal tract that were largely independent of anatomic location, suggesting the presence of shared biology across anatomical boundaries (Liu et al. 2018). Whereas these studies helped establish histology-wide similarities, our work defines the specific tumor-normal differential expression program underlying adenocarcinoma biology across organ systems and leverages this program for both drug discovery and patient prognostication. Heterogeneity in the identified transcriptomic profile is associated with patient outcomes, suggesting that the profile associated with high-risk patients may be used for additional molecularly guided treatment.

With the shared biology of adenocarcinomas established, we sought to identify drugs that target vulnerabilities across these cancers. By leveraging the shared features, drug development efforts may be accelerated compared to studies siloed by tumor organ of origin. Connectivity map analysis was performed and identified compounds predicted to reverse the adenocarcinoma transcriptional phenotype towards a normal-like state. The result of this initial drug screen were supported by the positive identification of the tyrosine kinase inhibitors crizotinib and gefitinib, both currently used to treat adenocarcinomas given recurrent ALK/ROS1 fusions, MET alterations, and EGFR pathway activation in many adenocarcinomas (Huang et al. 2020; Ou et al. 2012; Zan et al. 2025). Vismodegib, a smoothened receptor inhibitor, and losartan, a ACE2i and TGFβ modulator, both have been shown to alter desmoplastic tumor stroma, especially in pancreatic and gastric cancer. The COX2 and HMGCOA reductase inhibitors, rofecoxib and atorvastatin, are widely prescribed, and have been proposed as potentially efficacious adjuncts to traditional anti-cancer therapy. *COXII* overexpression has been observed in multiple adenocarcinomas including those arising in GI, lung, and pancreas. Enhanced COXII activities are associated with increased proliferation, migration, and the immune environment, whereas the tumors often become dependent on the mevalonate pathway for prenylation (Habib et al. 2007; Sobolewski et al. 2010). These identified drugs and classes represent prioritized candidates for further testing in adenocarcinomas given their association with the described molecular patterns and literature supporting their mechanistic function in these tumors.

Limitations of the study include reliance on bulk RNA-sequencing datasets that are publicly available. Bulk RNA-sequencing data do not fully resolve tumor intrinsic *vs*. microenvironmental changes, but may be more readily translated than alternative single cell approaches. Second, connectivity map predictions are computational and require downstream experimental validation. Finally, although the pan-adenocarcinoma program captures commonalities across lung, gastric, and colorectal adenocarcinomas, tissue-specific biology cannot be fully disregarded and may influence drug response. Future work should test the pan-adenocarcinoma program in additional tumor types, integrate dedicated pharmacogenomic analysis to develop predictive biomarkers, and validate candidate drugs and biomarkers in preclinical models.

## Funding

This study did not receive any funding.

## Conflict of Interest

Authors declare no conflict of interest.

## Acknowledgements

M.D. would like to thank Dr. D. Keskin and J. Duke-Cohan for suggestions on analysis methods; Z.Z. would like to thank Dr. H. Feng for guidance on generating publication quality figures.

## Author Contributions

M.D. conceptualized the study, conducted analyses, generated figures, and wrote the manuscript. Z.Z. conducted analyses, generated figures, and wrote the manuscript. C.L. conducted analyses, generated figures, and wrote the manuscript.

## Data Availability

All data produced in the present study are available upon reasonable request to the authors.

## Notes

### Competing Interest Statement

The authors have declared no competing interest.

## References

Aitken M. 2023. Global Oncology Trends-2023. The IQVIA Institute.

Biller LH, and Schrag D. 2021. Diagnosis and Treatment of Metastatic Colorectal Cancer: A Review. JAMA 325:669–685. 10.1001/jama.2021.0106

Bray F, Ferlay J, Soerjomataram I, Siegel RL, Torre LA, and Jemal A. 2018. Global cancer statistics 2018: GLOBOCAN estimates of incidence and mortality worldwide for 36 cancers in 185 countries. CA Cancer J Clin 68:394–424. 10.3322/caac.21492

Christenson ES, Jaffee E, and Azad NS. 2020. Current and emerging therapies for patients with advanced pancreatic ductal adenocarcinoma: a bright future. Lancet Oncol 21:e135–e145. 10.1016/S1470-2045(19)30795-8

Collaborators GUHD. 2023. The burden of stomach cancer mortality by county, race, and ethnicity in the USA, 2000-2019: a systematic analysis of health disparities. Lancet Reg Health Am 24:100547. 10.1016/j.lana.2023.100547

Didier AJ, Nandwani S, Fahoury AM, Craig DJ, Watkins D, Campbell A, Spencer CT, Batten M, Vijendra D, and Sutton JM. 2024. Trends in pancreatic cancer mortality in the United States 1999-2020: a CDC database population-based study. Cancer Causes Control 35:1509–1516. 10.1007/s10552-024-01906-z

Habib A, Shamseddeen I, Nasrallah MS, Antoun TA, Nemer G, Bertoglio J, Badreddine R, and Badr KF. 2007. Modulation of COX-2 expression by statins in human monocytic cells. FASEB J 21:1665–1674. 10.1096/fj.06-6766com

Hoadley KA, Yau C, Hinoue T, Wolf DM, Lazar AJ, Drill E, Shen R, Taylor AM, Cherniack AD, Thorsson V, Akbani R, Bowlby R, Wong CK, Wiznerowicz M, Sanchez-Vega F, Robertson AG, Schneider BG, Lawrence MS, Noushmehr H, Malta TM, Cancer Genome Atlas N, Stuart JM, Benz CC, and Laird PW. 2018. Cell-of-Origin Patterns Dominate the Molecular Classification of 10,000 Tumors from 33 Types of Cancer. Cell 173:291–304 e296. 10.1016/j.cell.2018.03.022

Huang L, Jiang S, and Shi Y. 2020. Tyrosine kinase inhibitors for solid tumors in the past 20 years (2001-2020). J Hematol Oncol 13:143. 10.1186/s13045-020-00977-0

Jeon H, Wang S, Song J, Gill H, and Cheng H. 2025. Update 2025: Management of Non-Small-Cell Lung Cancer. Lung 203:53. 10.1007/s00408-025-00801-x

Joshi SS, and Badgwell BD. 2021. Current treatment and recent progress in gastric cancer. CA Cancer J Clin 71:264–279. 10.3322/caac.21657

Kratzer TB, Bandi P, Freedman ND, Smith RA, Travis WD, Jemal A, and Siegel RL. 2024. Lung cancer statistics, 2023. Cancer 130:1330–1348. 10.1002/cncr.35128

Lamb J, Crawford ED, Peck D, Modell JW, Blat IC, Wrobel MJ, Lerner J, Brunet JP, Subramanian A, Ross KN, Reich M, Hieronymus H, Wei G, Armstrong SA, Haggarty SJ, Clemons PA, Wei R, Carr SA, Lander ES, and Golub TR. 2006. The Connectivity Map: using gene-expression signatures to connect small molecules, genes, and disease. Science 313:1929–1935. 10.1126/science.1132939

Lin EW, Karakasheva TA, Lee DJ, Lee JS, Long Q, Bass AJ, Wong KK, and Rustgi AK. 2017. Comparative transcriptomes of adenocarcinomas and squamous cell carcinomas reveal molecular similarities that span classical anatomic boundaries. PLoS Genet 13:e1006938. 10.1371/journal.pgen.1006938

Liu Y, Sethi NS, Hinoue T, Schneider BG, Cherniack AD, Sanchez-Vega F, Seoane JA, Farshidfar F, Bowlby R, Islam M, Kim J, Chatila W, Akbani R, Kanchi RS, Rabkin CS, Willis JE, Wang KK, McCall SJ, Mishra L, Ojesina AI, Bullman S, Pedamallu CS, Lazar AJ, Sakai R, Cancer Genome Atlas Research N, Thorsson V, Bass AJ, and Laird PW. 2018. Comparative Molecular Analysis of Gastrointestinal Adenocarcinomas. Cancer Cell 33:721–735 e728. 10.1016/j.ccell.2018.03.010

Love MI, Huber W, and Anders S. 2014. Moderated estimation of fold change and dispersion for RNA-seq data with DESeq2. Genome Biol 15:550. 10.1186/s13059-014-0550-8

Ou SH, Bartlett CH, Mino-Kenudson M, Cui J, and Iafrate AJ. 2012. Crizotinib for the treatment of ALK-rearranged non-small cell lung cancer: a success story to usher in the second decade of molecular targeted therapy in oncology. Oncologist 17:1351–1375. 10.1634/theoncologist.2012-0311

Schumacher BLaM. 1992. Maximally Selected Rank Statistics. Biometrics 48:73–85.

Siegel RL, Miller KD, Wagle NS, and Jemal A. 2023. Cancer statistics, 2023. CA Cancer J Clin 73:17–48. 10.3322/caac.21763

Sinicrope FA. 2022. Increasing Incidence of Early-Onset Colorectal Cancer. N Engl J Med 386:1547–1558. 10.1056/NEJMra2200869

Sobolewski C, Cerella C, Dicato M, Ghibelli L, and Diederich M. 2010. The role of cyclooxygenase-2 in cell proliferation and cell death in human malignancies. Int J Cell Biol 2010:215158. 10.1155/2010/215158

Subramanian A, Narayan R, Corsello SM, Peck DD, Natoli TE, Lu X, Gould J, Davis JF, Tubelli AA, Asiedu JK, Lahr DL, Hirschman JE, Liu Z, Donahue M, Julian B, Khan M, Wadden D, Smith IC, Lam D, Liberzon A, Toder C, Bagul M, Orzechowski M, Enache OM, Piccioni F, Johnson SA, Lyons NJ, Berger AH, Shamji AF, Brooks AN, Vrcic A, Flynn C, Rosains J, Takeda DY, Hu R, Davison D, Lamb J, Ardlie K, Hogstrom L, Greenside P, Gray NS, Clemons PA, Silver S, Wu X, Zhao WN, Read-Button W, Wu X, Haggarty SJ, Ronco LV, Boehm JS, Schreiber SL, Doench JG, Bittker JA, Root DE, Wong B, and Golub TR. 2017. A Next Generation Connectivity Map: L1000 Platform and the First 1,000,000 Profiles. Cell 171:1437–1452 e1417. 10.1016/j.cell.2017.10.049

Sun D, Gao, W., Hu, H., and Zhou, S. 2022. Why 90% of clinical drug development fails and how to improve it? . Acta Pharm Sin B12:3049–3062.

Sung H, Ferlay J, Siegel RL, Laversanne M, Soerjomataram I, Jemal A, and Bray F. 2021. Global Cancer Statistics 2020: GLOBOCAN Estimates of Incidence and Mortality Worldwide for 36 Cancers in 185 Countries. CA Cancer J Clin 71:209–249. 10.3322/caac.21660

Wu T, Hu E, Xu S, Chen M, Guo P, Dai Z, Feng T, Zhou L, Tang W, Zhan L, Fu X, Liu S, Bo X, and Yu G. 2021. clusterProfiler 4.0: A universal enrichment tool for interpreting omics data. Innovation (Camb) 2:100141. 10.1016/j.xinn.2021.100141

Yoav Benjamini YH. 1995. Controlling the False Discovery Rate: A Practical and Powerful Approach to Multiple Testing. Journal of the Royal Statistical Society: Series B (Methodological) 57:289–300.

Yu G, Wang LG, Han Y, and He QY. 2012. clusterProfiler: an R package for comparing biological themes among gene clusters. OMICS 16:284–287. 10.1089/omi.2011.0118

Zan H, Xu M, Guo P, and Yu X. 2025. Efficacy and safety of EGFR-TKI for EGFR-mutated NSCLC: systematic review and network meta-analysis. Int J Clin Exp Pathol 18:386–404. 10.62347/DWIW6941

