## Supplementary Information for "A Shared Adenocarcinoma Transcriptomic Program Enables Prediction of Therapeutics Applicable Across Tissues"

**Supplementary Figure 1: Dimensional reduction of tumor *vs.* normal tissue samples by PCA.** (A–C) PCA plots of tumor (orange) vs. normal (green) samples from LUAD (A), STAD (B), and COAD (C) patients. LUAD and COAD exhibit clear group separation, while STAD shows moderate overlap with its matched normal tissue samples.

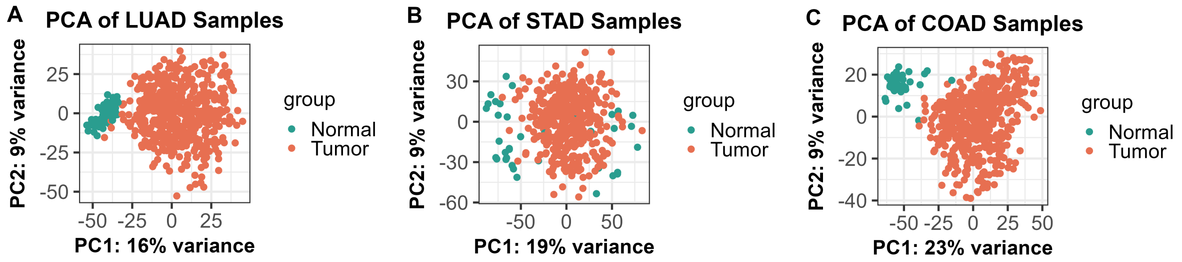


**Supplementary Figure 2: Mean-normalized visualization of DEGs by MA plots.** (A–B) MA plots showing $log_2$ fold changes vs. mean-normalized counts for LUAD (A), STAD (B), and COAD (C), compared to its matched normal tissue samples. Blue points represent significantly differentially expressed genes.

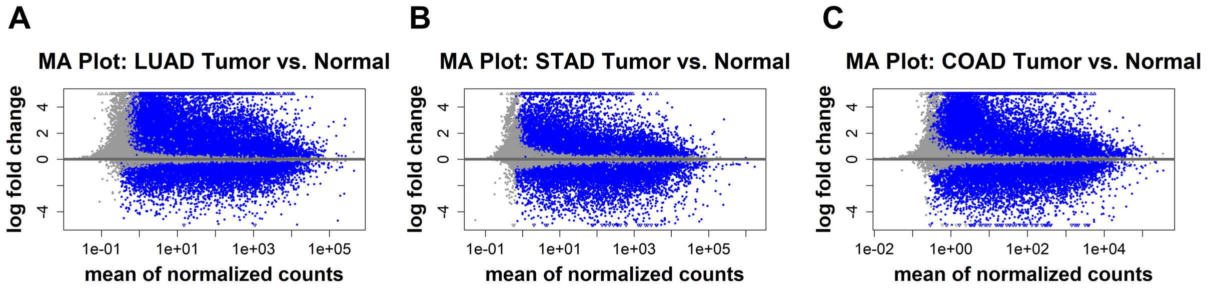

**Supplementary Figure 3: UpSet plot visualization of consensus DEGs.** UpSet plots show intersecting sets of upregulated (A) and downregulated (B) genes between different comparison groups.

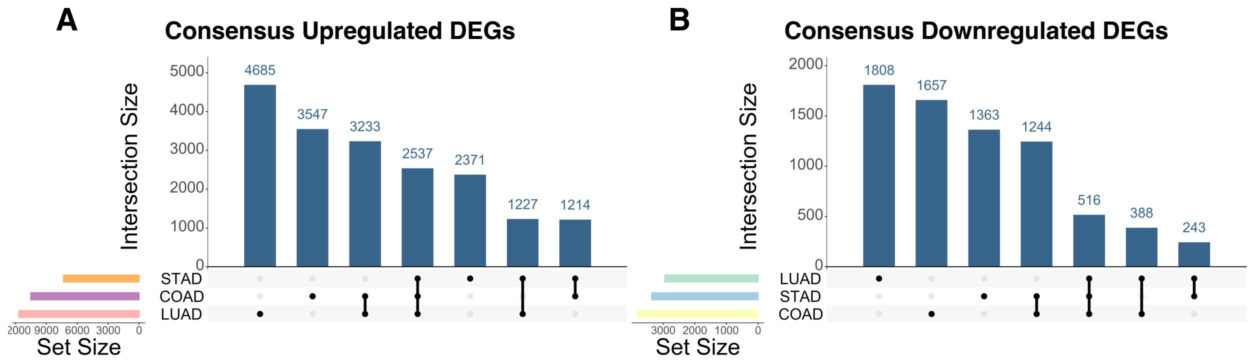

**Supplementary Figure 4: The consensus DEG signature can predict the survival of patients with any of the three adenocarcinomas.** (A-C) Kaplan–Meier survival analysis of overall survival of patients with LUAD (A), STAD (B), or COAD (C), segregated by the expression of all of the consensus upregulated (top) or downregulated (bottom) DEGs shared by all three adenocarcinomas. Patients were stratified by optimal cutoff. Statistical analysis was conducted by the log-rank test.

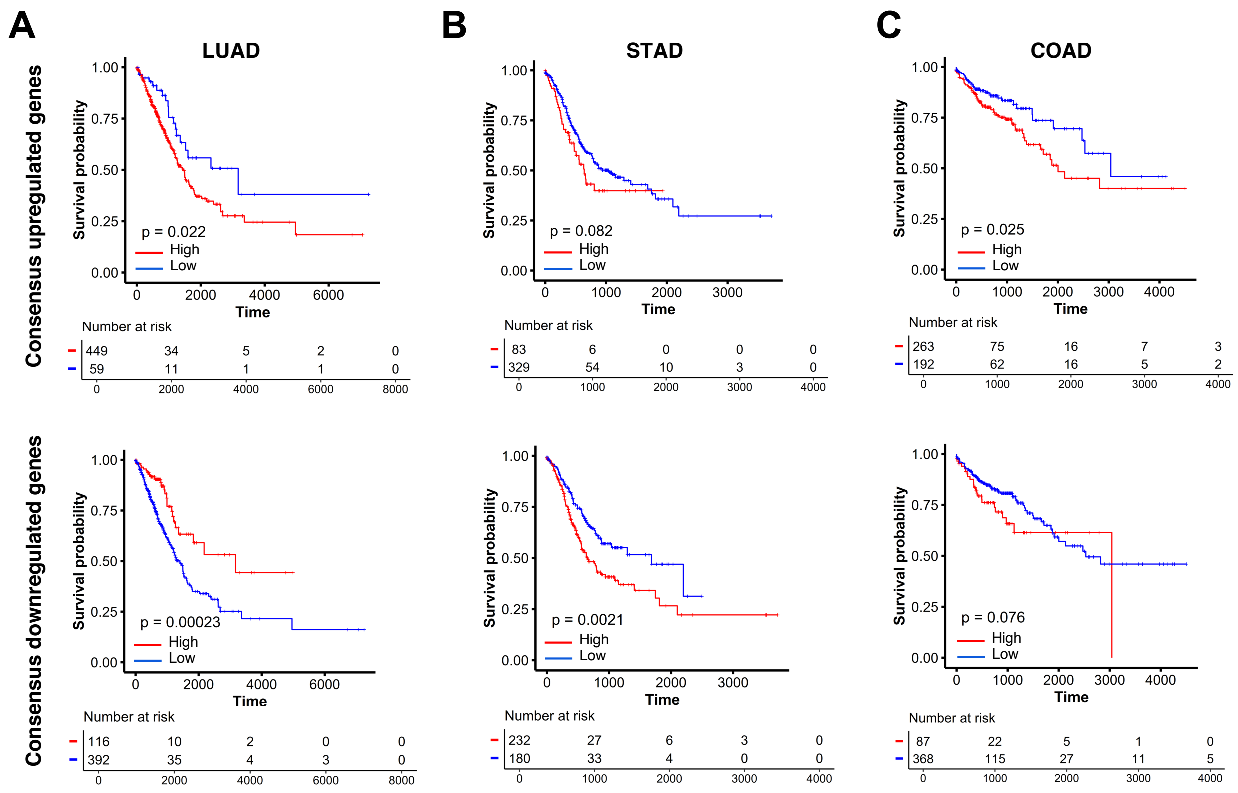

**Supplementary Figure 5: Consensus DEG clusters show a trend of predicting STAD patient survival.** (A) Heatmap of unsupervised hierarchical clustering of the 3053 consensus DEGs shared by all three adenocarcinomas categorize STAD patients into two subgroups. (B) Kaplan-Meier plot of the two primary patient clusters shows a trend in segregating patient survival (p = 0.13). Statistical analysis was conducted by the log-rank test.

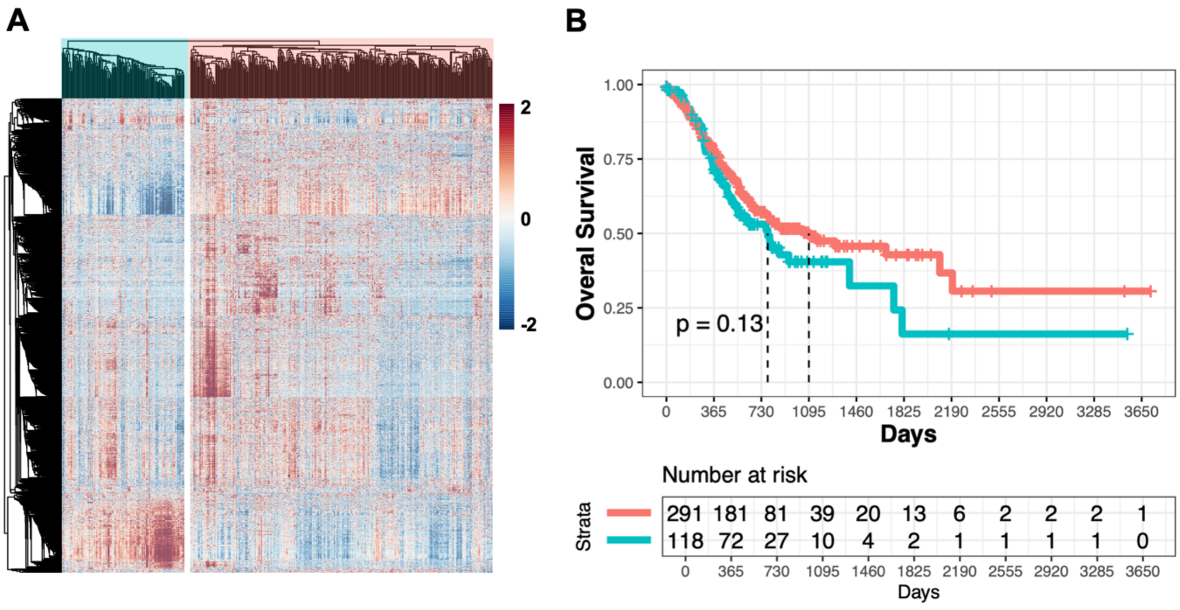

**Supplementary Figure 6: Consensus DEG clusters show a trend of predicting STAD patient survival.** (A) Heatmap of unsupervised hierarchical clustering of the 3053 consensus DEGs shared by all three adenocarcinomas categorize STAD patients into three subgroups. (B) Kaplan-Meier plot of the three primary patient clusters shows a trend in segregating patient survival (p=0.3). Statistical analysis was conducted by the log-rank test.

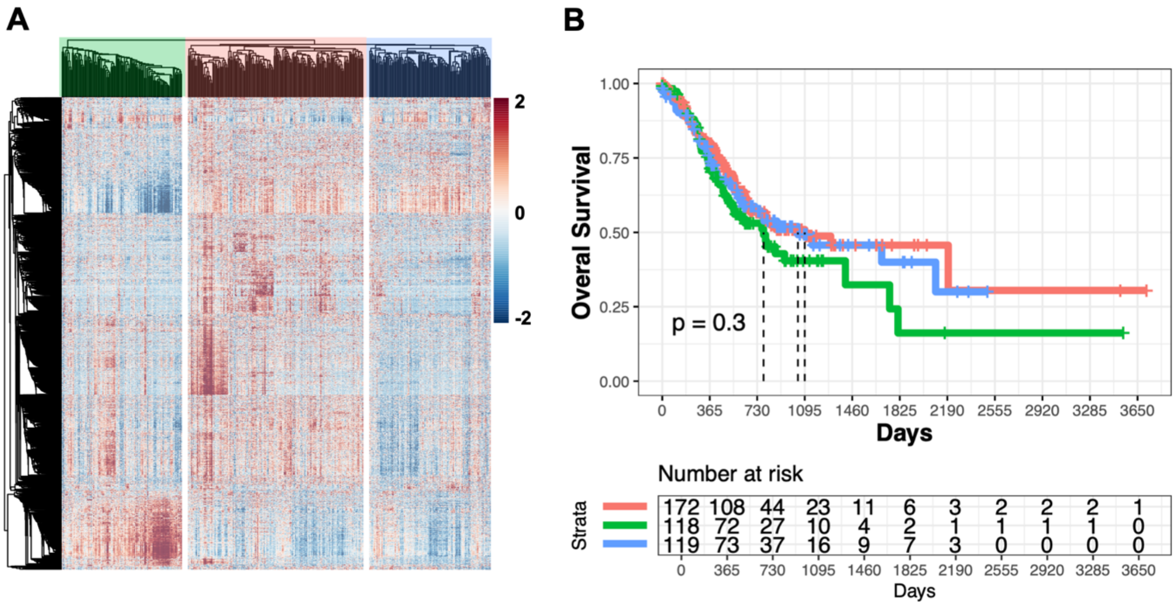

**Supplementary Figure 7: Consensus DEG clusters show a trend of predicting COAD patient survival.** (A) Heatmap of unsupervised hierarchical clustering of the 3053 consensus DEGs shared by all three adenocarcinomas categorize COAD patients into two subgroups. (B) Kaplan-Meier plot of the two primary patient clusters shows a trend in segregating patient survival (p = 0.21). Statistical analysis was conducted by the log-rank test.

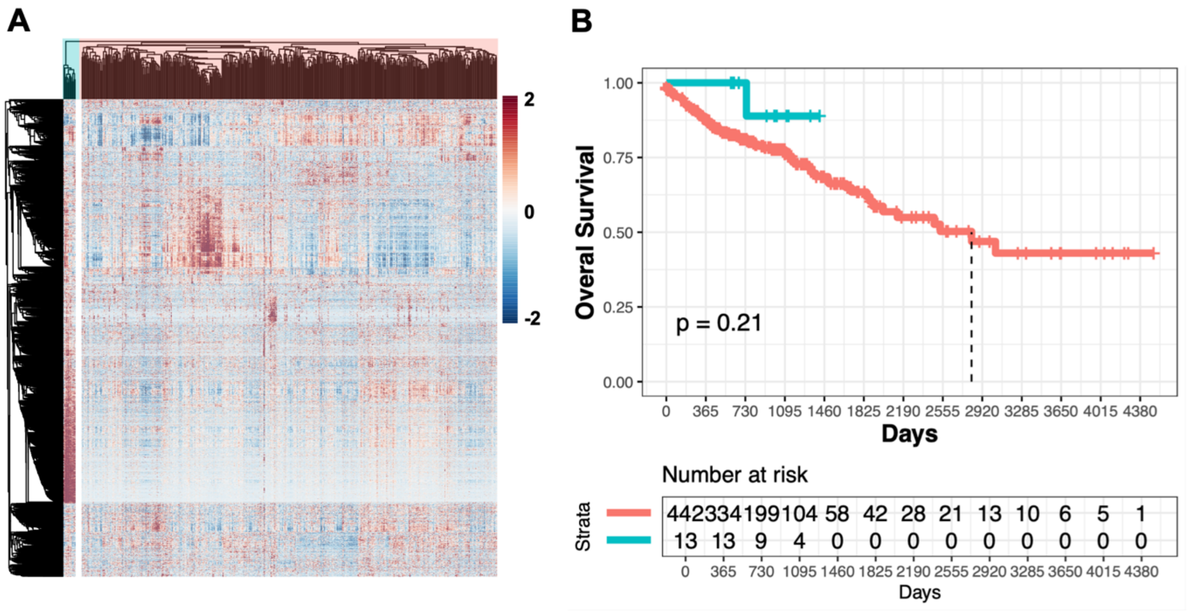

**Supplementary Figure 8: Consensus DEG clusters show a trend of predicting COAD patient survival.** (A) Heatmap of unsupervised hierarchical clustering of the 3053 consensus DEGs shared by all three adenocarcinomas categorize COAD patients into three subgroups. (B) Kaplan-Meier plot of the three primary patient clusters shows a trend in segregating patient survival (p = 0.25). Statistical analysis was conducted by the log-rank test.

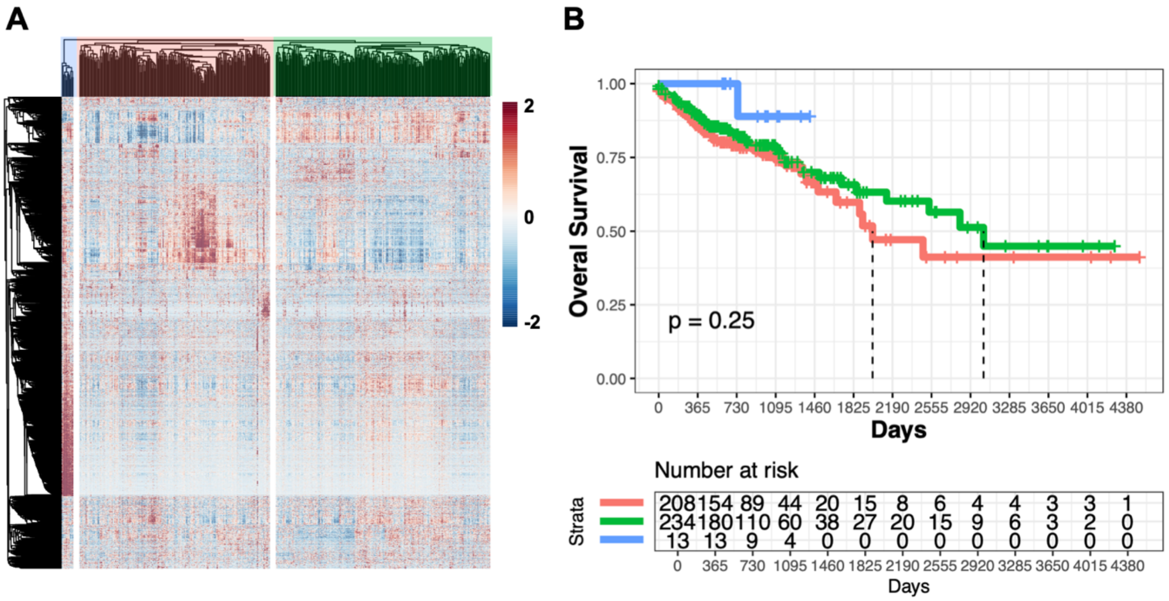
